# Adhesive defence mucus secretions in the red triangle slug (*Triboniophorus graeffei*) can incapacitate adult frogs

**DOI:** 10.1101/544775

**Authors:** John Gould, Jose W. Valdez, Rose Upton

**Affiliations:** School of Environmental and Life Sciences, University of Newcastle, Callaghan, 2308 NSW, Australia; Department of Bioscience - Biodiversity and Conservation, Aarhus University, 8410 Rønde, Denmark

**Keywords:** antipredator, adhesive gel, bioadhesive, *Litoria chloris*, mollusc, predator-prey interactions

## Abstract

Gastropods are known to secrete mucus for a variety of purposes, including locomotion, reproduction, adhesion to surfaces, and lubrication. A less commonly known function of mucus secretion in this group involves its use as a defence against predation. Among the terrestrial slugs, mucus that serves this particular purpose has been studied for only a handful of species under laboratory conditions, where it is thought to be produced for self-fouling or to make individuals difficult to consume. However, the mechanisms of how these defensive secretions operate and their effectiveness in deterring predation in the natural world have not be described in much detail. In this study, we provide evidence of adhesive mucus secretions in the red triangle slug (*Triboniophorus graeffei*) as an adaptation against predation. Field observations of a large red-eyed green tree frog (*Litoria chloris*) trapped in the mucus secretions of a nearby *T. graeffei* revealed that this mucus serves to incapacitate predators rather than just simply as an overall deterrence. Mechanical stimulation of *T. graeffei* under laboratory conditions revealed that adhesive secretions were produced from discrete sections of the dorsal surface when disturbed, leading to the production of a highly sticky and elastic mucus that was unlike the thin and slippery mucus used during locomotion. The adhesiveness of the defensive secretions was strengthened and reactivated when in contact with water. This appears to not only be the first description of defensive mucus production in this slug species but one of the first natural observations of the use of adhesive defence secretions to incapacitate a predator. The biomechanical properties of this mucus and its ability to maintain and strengthen its hold under wet conditions make it potentially useful in the development of new adhesive materials.

## Introduction

Animals have evolved a diverse array of anti-predator traits, including those that are physical (camouflage, mimicry, and weaponry), behavioural (defensive displays, colouration), and chemical (venom, noxious chemicals). The production of mucus is a prime example of a chemical response to predation risk which has been recorded amongst velvet worms (Baer & Mayer, 2012), echinoderms (Flammang, Demeuldre, Hennebert, & Santos, 2016), fish (Schubert, Munday, Caley, Jones, & Llewellyn, 2003; Shephard, 1994), arthropods (Betz & Kölsch, 2004), lizards (Brau, Lanterbecq, Zghikh, Bels, & Damman, 2016), aquatic gastropods (Rice, 1985), terrestrial slugs (Barber et al., 2015; Deyrup-Olsen, Luchtel, & Martin, 1983), and amphibians (Arnold, 1982; Evans & Brodie, 1994; Graham, Glattauer, Li, Tyler, & Ramshaw, 2013). Such bioadhesives are typically secreted quickly and exhibit a rapid curing process, with some able to be exposed for weeks without losing their bonding capability (von Byern et al., 2017). These secretions are produced at the onset of an attack, most noticeably mechanical stimulation, acting as a protective barrier or overall deterrent that reduces the chances of the prey species from being consumed.

Gastropods are the archetypal animals known for their viscoelastic mucous secretions which aid in locomotion, reproduction, and adhesion to surface substrates while foraging (Smith, 2010). Many gastropod species which have transitioned to a life on land also secrete a thin coating of mucosa in order to remain well lubricated, as their soft bodies and permeable epidermal linings mean they are particularly vulnerable to mechanical damage and desiccation (South, 2012; Verdugo, 1991). In addition to these well-known functions of mucus secretions amongst the gastropods, some species have also evolved supplementary secretions that serve to reduce the threat of predation (Rollo & Wellington, 1979; Triebskorn & Ebert, 1989). This has been recorded among the terrestrial slugs, which are very susceptible to predation due to their lack of a protective shell, soft bodies and slow pace. This defence-specific mucus is often distinct from the usually thin and slippery mucus produced for the purposes of lubrication and locomotion, acting as a protective barrier that prevents contact between the potential threat and the slug’s body (Deyrup-Olsen et al., 1983; Smith, 2006).

Although defensive mucus secretions in terrestrial slugs are thought to serve as methods of self-fouling that deters predation, recent findings have found that some species secrete mucus with adhesive properties as part of their anti-predation repertoire (Foltan, 2004; Landauer & Chapnick, 1981; Rice, 1985; Smith, 2010). For example, *Arion subfuscus* and *Ariolimax columbianus* have been shown to possess dorsal epithelial that secretes an adhesive defensive mucus when disturbed (Deyrup-Olsen et al., 1983; Mair & Port, 2002; Martin & Deyrup-Olsen, 1986). In these species, the defensive mucus starts out as a viscous slime, which sets into a highly sticky and elastic mass which, although composed of 95% water, can sustain stressor over 100 kPa due to gel-stiffening proteins which bind with metals (Smith, 2010; Wilks, Rabice, Garbacz, Harro, & Smith, 2015). Likewise, slugs in the genera *Veronicella* also produce their own form of sticky mucus in response to irritation (Cook, 1987). The mechanical properties of these adhesive secretions are so remarkable they have inspired the development of new surgical glues (Li et al., 2017), and are thought to confer an evolutionary advantage for terrestrial slugs by making them unpalatable or difficult to consume.

The use of adhesive mucus across multiple genera suggests that it may be a common defence mechanism among gastropods, particularly terrestrial slugs which lack a protective structure. However, studies on defensive secretions in gastropods have been conducted nearly exclusively on the terrestrial slugs *A. subfuscus* and *A. columbianus* (Smith & Callow, 2006). This presents an issue since there are likely to be many more species with defensive mucus secretions that have unique adhesive properties due to differences in the composition of their mucus secretions (Foltan, 2004). Moreover, many of these studies test the properties of adhesive secretions in a laboratory setting where their use as a form of antipredator defence is assumed. As such, the mechanism of how adhesive defence mucus operates in the natural world and their effectiveness in deterring predation has not been described in much detail in the literature to date. In this study, we show the first evidence of adhesive mucus secretions in the red triangle slug (*Triboniophorus graeffei*) as an adaptive response to predation threat. We also show that these secretions can act to incapacitate predators rather than simply methods of self-fouling that make individuals unpalatable or noxious to predators.

## Materials and Methods

Field observations occurred within the Watagans Mountain Range, New South Wales, Australia on October 27, 2017. Nightly fieldwork resulted in the chance discovery of an adult male red-eyed green tree frog (*Litoria chloris*) that appeared to be stuck to a fallen eucalyptus branch during a period of heavy rainfall. Since the frog was found in close proximity to a large *T. graeffei* individual, it was hypothesised that it had become stuck due to the secretions of the slug, either by misfortune or after a predation attempt. Observations were made in the field for a period of ten minutes without interference to examine any change in the situation and whether *L. chloris* would free itself. No changes occurred so the frog and the nearby slug were then taken back to the Conservation Biology laboratory at the University of Newcastle for further investigation.

The adult frog, while still attached to the branch, was transferred to a 27 x 17 x 15 cm container will with 2-3 cm of aged tap water and regularly checked for recovery or signs of stress until it was no longer adhered to the branch. The *T. graeffei* individual was placed onto a petri dish, where the typical mucus properties exhibited during locomotion were examined. To determine whether it produced adhesive secretions that could explain the incapacitated state of the adult frog, mucus production was encouraged by disturbance via mechanical stimulation for a period of 60 seconds using a gloved finger. On November 24, 2018, an additional three *T. graeffei* individuals were collected from the Watagans Mountain Range to gain additional information regarding the potential adhesive properties of this defensive mucus. Specimens were housed together in a 27 x 17 x 15 cm container with leaf litter for seven days. Over this period, we evaluated the quantity, thickness, and adhesive quality of the mucus left behind during locomotion, as well as the mucus of the dorsal surface before and after 60 seconds of tactile stimulation.

## Results

Observations made in the field indicate that the ventral skin area of *L. chloris* was strongly adhered to the surface of the branch, including the lower throat, abdomen, and inner thighs of the hind legs (Fig. 1). Particularly noteworthy was the unusual positioning of the frog, with the body very close to the surface of the branch and the legs splayed out (Fig. 2). Additionally, the toe pads and webbing of the front legs were bonded to each other and partially to the branch, while the toe pads of the back legs were also stuck to the branch (Fig. 2). On multiple occasions during field observations, the individual attempted escape but was unable to remove itself from its unorthodox position. Attempts to physically remove the frog also failed, with the individual producing a distress call each time. Closer observations of the branch and surrounding leaf litter showed no signs that the frog was covered in fallen sap from nearby trees. Throughout this period of field observations, the *T. graeffei* individual did not move, remaining less than 1 cm away from the mouth of the frog (Fig. 1, Fig. 2).

**Figure 1.**
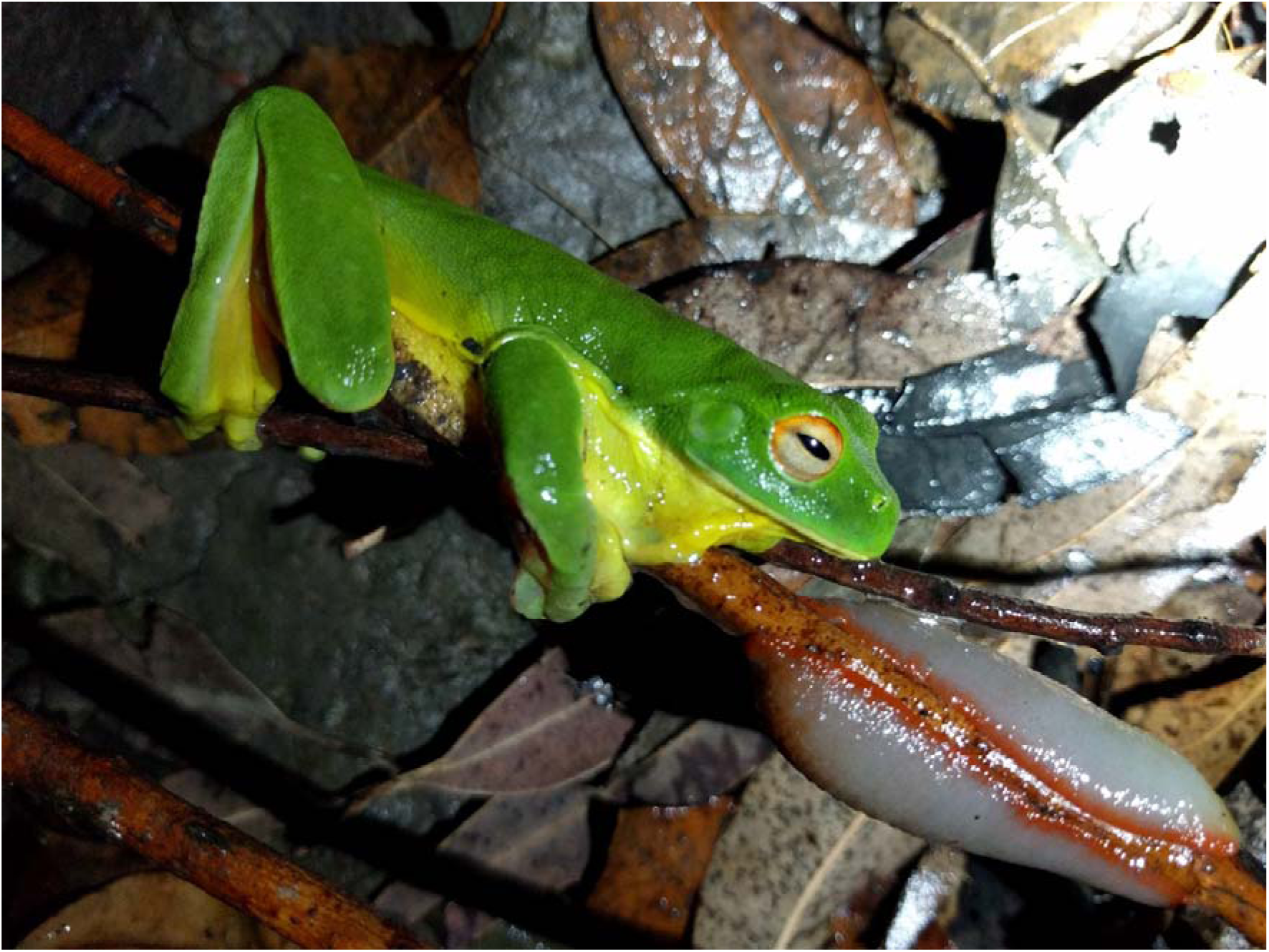
Adult *Litoria chloris* adhered to a eucalyptus branch in close proximity to a *Triboniophorus graeffei* individual. The frog is in an unusual position, with the skin of the throat stretched.

**Figure 2.**
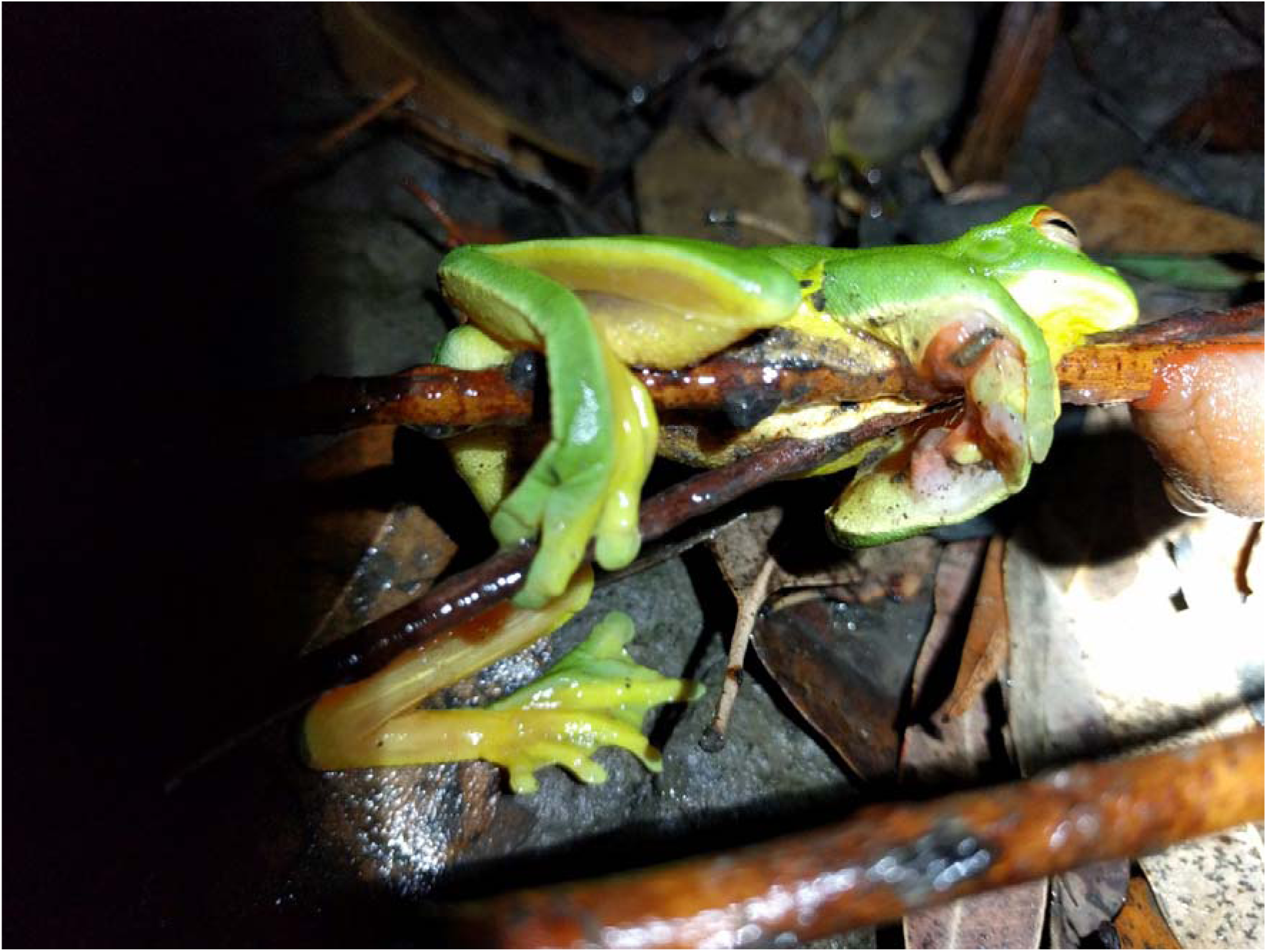
Adult *Litoria chloris* adhered to a eucalyptus branch with the front toes fixed to each other around the branch.

Following its collection, the frog remained attached to the branch, for a period of 48 hours with the skin of the abdomen, legs, and toes remaining covered in the sticky residue. until the skin was sloughed naturally. Additional residue was also present on the section of the branch that was collected which, along with affected areas of skin, appeared to be mostly translucent with sections that were slightly red in colouration (Fig. 3). After more than 24 hours it was still unable to remove itself, so to prevent any further distress we assisted in its removal by carefully peeling away sections of skin that were adhered to the branch. Even after the frog was no longer stuck to the branch, its skin remained covered in mucus that would often cause it to become adhered to the bottom of the container while it was immersed in water.

**Figure 3.**
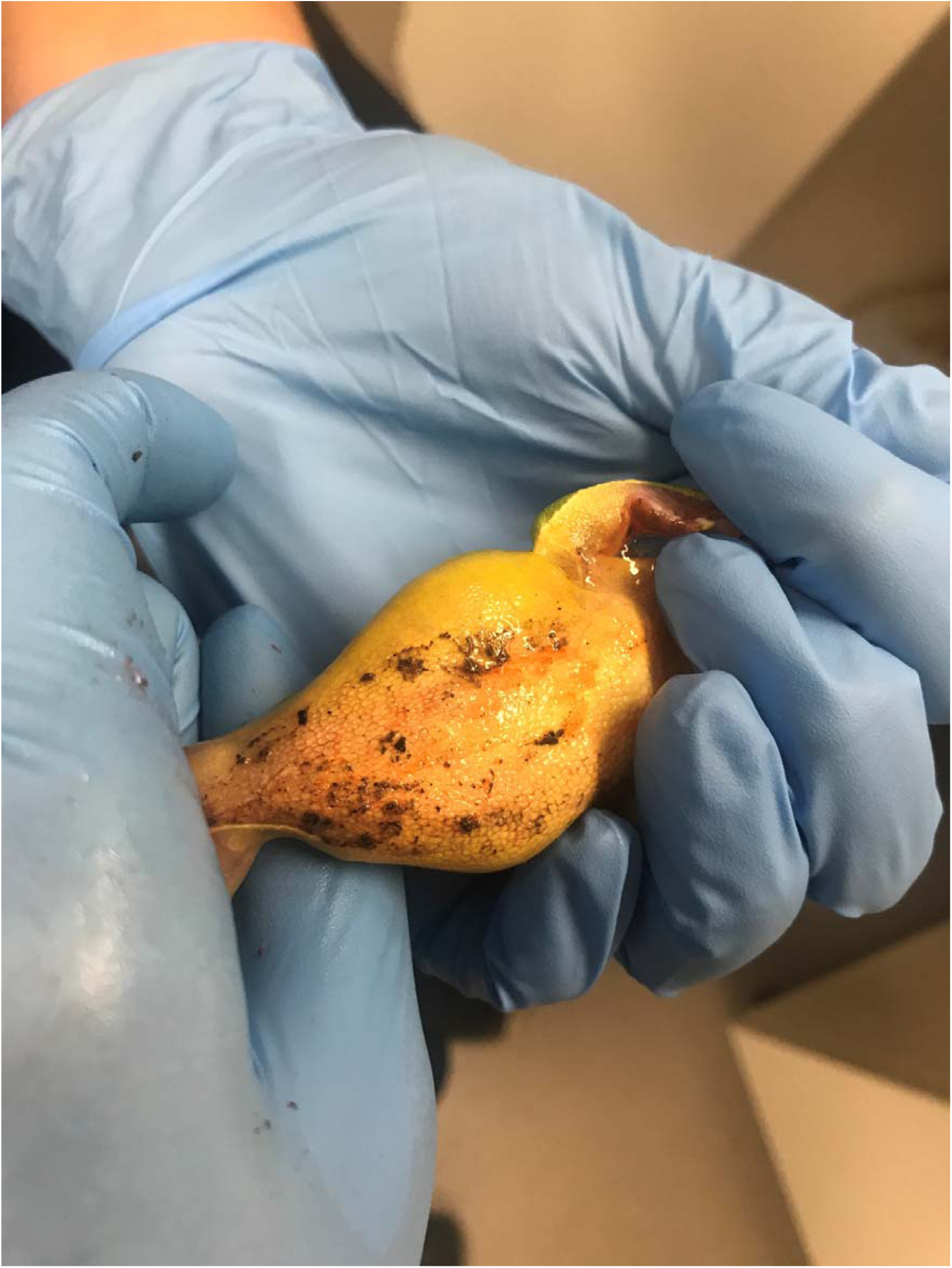
Underside of adult *Litoria chloris* covered with adhesive mucus and slight red coloration.

When examining the *T. graeffei* individual, the mucus layer left behind during periods of locomotion was found to be quite thin and lacking in adhesiveness (Fig. 4). Following the examination of the additional specimens that were collected, it was discovered that the dorsal surface was typically dry to the touch prior to disturbance but became wet with copious amounts of extremely adhesive mucus during periods of mechanical stimulation. Only those portions of dorsum that were stimulated showed an increased expression of mucus while surrounding regions remained relatively dry. Often a single touch of the dorsum resulted in contractions in the area and the immediate secretion of mucus (Fig. 5), which was expelled onto the surface in the form of tiny droplets that quickly spread over the surrounding surface. This mucus became adhesive within a matter of seconds and often resulted in the fingers of gloved hands becoming stuck together or to surrounding paper and plastic when handled. However, the mucus gradually lost its adhesive quality over a few minutes, depending on quantity, as it began to desiccate, which was only gained upon rehydration. Once expressed, this mucus became increasingly thick, sticky and opaque with repeated stimulation. On some occasions, disturbance also resulted in the production of red coloured mucus which became dispersed throughout the mostly clear mucus. This particular mucus was only expressed during periods when mechanical stimulation was applied on the red tissue located on the perimeter around the foot of the slug (Fig. 6) and was similar to the residue present on the frog’s skin (Fig. 3).

**Figure 4.**
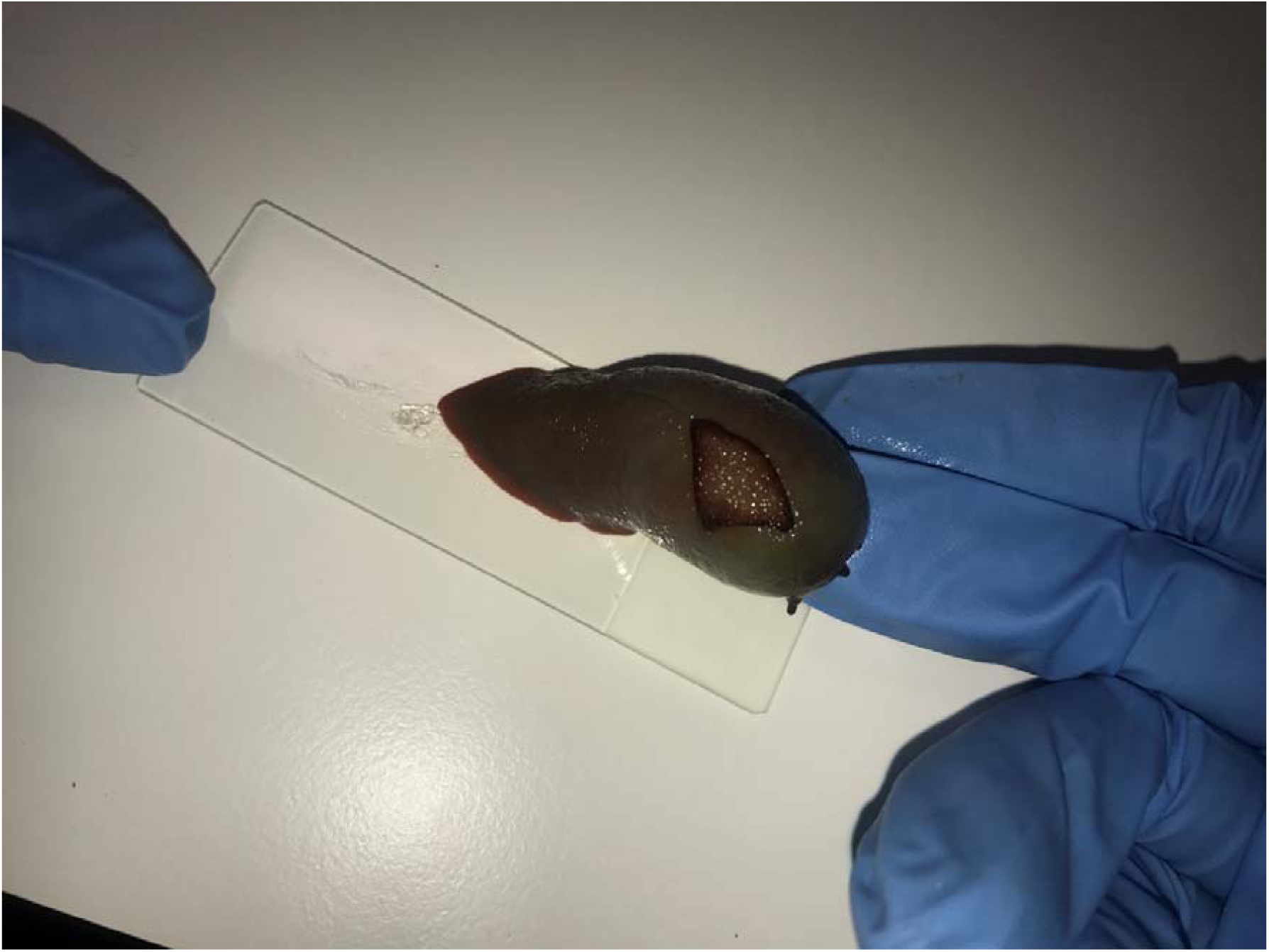
Analysis of mucus secretions used for locomotion from the base of the foot of *Triboniophorus graeffei*.

**Figure 5.**
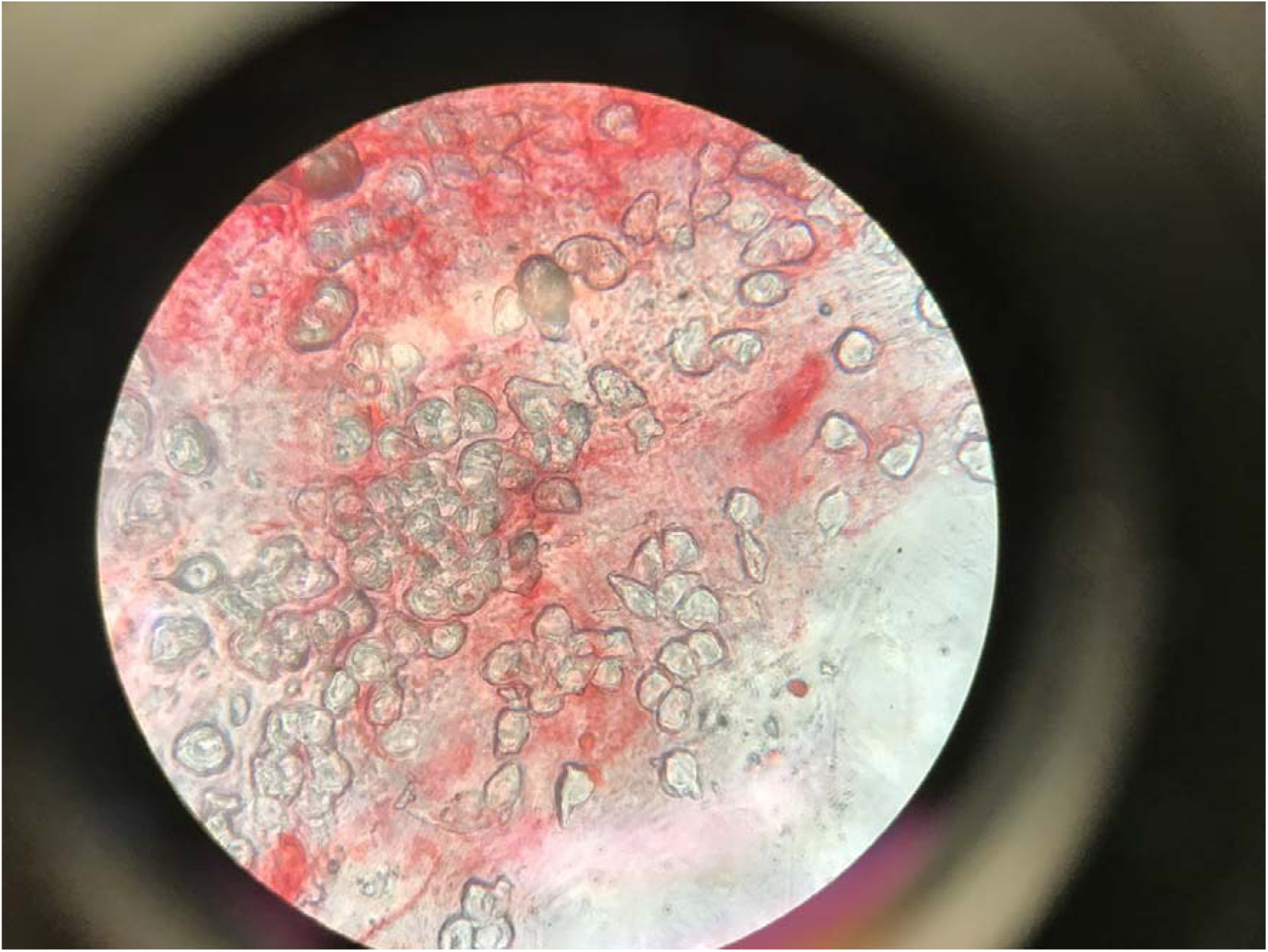
Microscopic photo of the mucus secretions from the dorsal surface of *Triboniophorus graeffei.*

**Figure 6.**
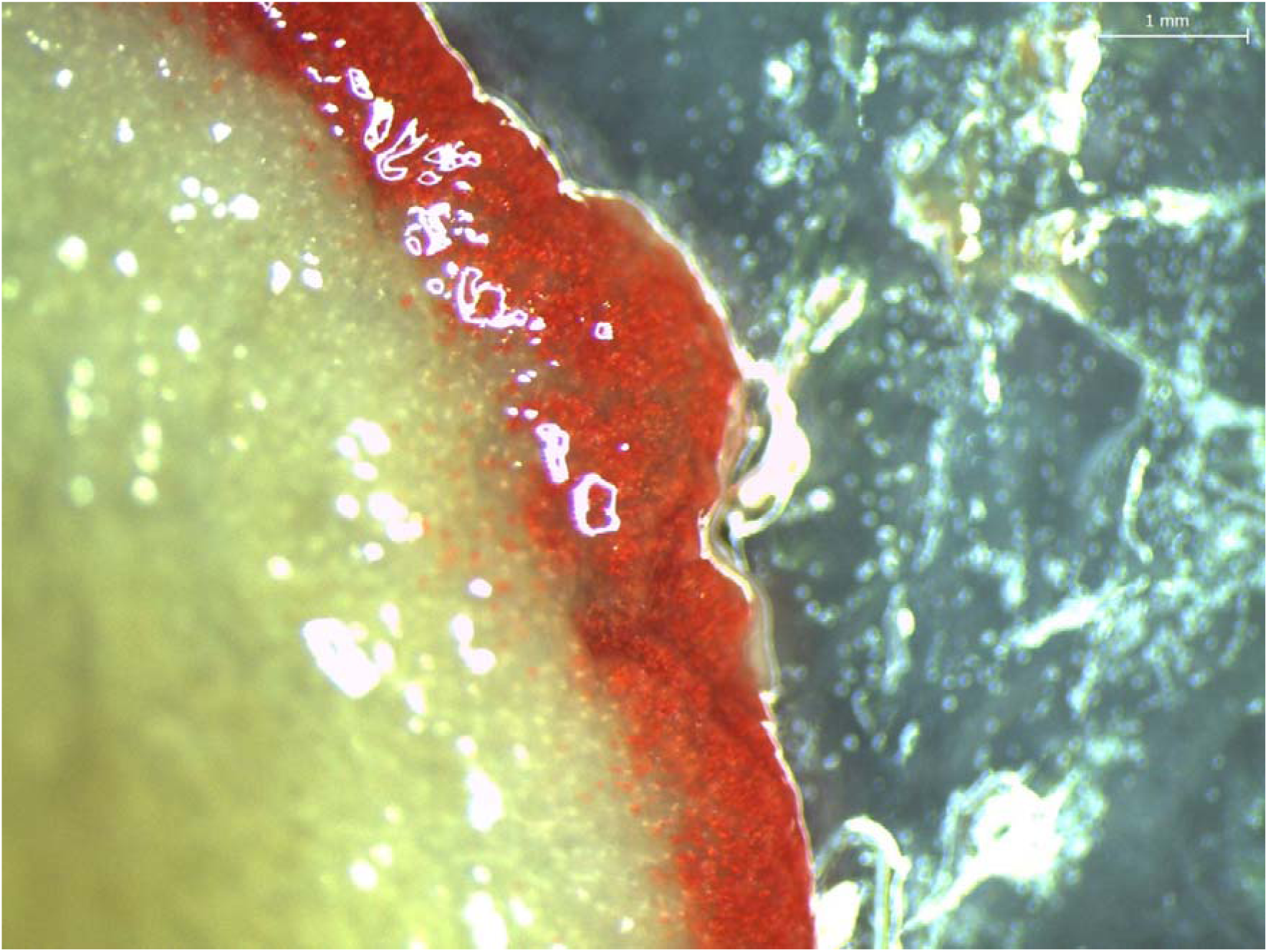
Red tissue located around the perimeter of the foot of *Triboniophorus graeffei* which produced a red coloured mucus which disturbed.

## Discussion

Our study indicates that *T. graeffei* secretes a highly adhesive mucus as an anti-predatory response that is distinct from the mucus produced for lubrication and locomotion. This defensive mucus has the ability to incapacitate comparatively large predators, such as frogs, for an extended period of time, which may facilitate escape or reduce the chance of a predator mounting further attacks while an escape is being made. The covering of *L. chloris* ventral surface in a sticky residue and its proximity to the *T. graeffei* specimen suggests that the frog activated an anti-predator response in the slug, resulting in it becoming covered in the adhesive mucus. This is also supported by the similarity in colouration of the mucus present on the frog and the red mucus often produced by *T. graeffei* when disturbed. The reddish colouration and the large quantity of mucus present on the frog’s skin, as well as the underlying branch, are in contrast to the thin and slippery locomotive mucus produced by the sole of the slug during locomotion, suggesting that the frog wasn’t simply trapped behind the slug by chance. Instead, it is more likely that the frog mounted a predatory attack on the slug, leading to the secretion of a large quantity of defensive mucus that resulted in it becoming stuck to the branch shortly after.

The slug found in proximity to the frog was able to produce sufficient mucus to incapacitate its frog predator in a state that would have likely persisted for at least two days, making the frog vulnerable to desiccation and its own predation. In order to incapacitate such a large predator, *T. graeffei* would have had to produce a large supply of mucus that could become adhesive in a relatively short period of time. The presence of mucus across nearly the entire ventral surface of the *L. chloris* adult and its close proximity to the slug are highly suggestive of these properties and further supported by analysis of the mucus under laboratory conditions. These qualities correspond with the defensive secretions produced by other slugs, such as *Ariolimax columbianus* and *Ario subfuscus*, which can produce copious amounts of mucus (5.5% of total body weight) that becomes adhesive within a matter of seconds (Deyrup-Olsen et al., 1983; Mair & Port, 2002; Martin & Deyrup-Olsen, 1986). The observations made in this study also demonstrate that the adhesive nature of the *T. graeffei* defence mucus would be reactivated repeatedly upon rehydration. Since amphibians have mucus glands to keep their skin moist, is likely the frog’s own secretions facilitated the sustained adhesiveness of the mucus, as well as its transference over a large surface area of its skin, especially if it began to struggle or attempted to remove the secretions using its foot pads. This may also have been further exacerbated by rainfall during this period, which may have sustained the adhesive quality of the mucus, while also allowing it to become more easily spread across the surface. Another possibility is that *L. chloris* could have produced its own adhesive secretions during its distress, which may have increased the adhesive properties of the slug mucus even further. Amphibians are known to release antipredator skin secretions which can be toxic or noxious, but many also species secrete adhesive substances which can be five times stronger than rubber cement (Evans & Brodie, 1994), and one experimental study on salamanders found that its adhesive mucus could incapacitate a snake predator for up to 48 hours (Arnold, 1982). Whether or not this was the case, other species also possess adhesive secretions for predation and the synergistic effect of such predator-prey adhesive secretions is not fully known and warrants further investigation.

Based on the experiments conducted in this study, it can be deduced that the cells responsible for secreting adhesive mucus in *T. graeffei* are located across the dorsal surface, which appear to be selectively activated in each discrete section of dorsum that becomes disturbed. Although many other regions have the capacity to produce mucus in this species, including the head, pneumostome, and sole, these do not seem to be involved in the production of defensive mucus. The exact mechanisms of mucus production in this species are yet to be determined, but it seems to be similar to production in *A. columbianus*, which also secrete adhesive mucus from glands located across the dorsal surface that is in contrast to the thin, slippery mucus secreted by the pedal foot (Luchtel, Deyrup-Olsen, & Martin, 1991; Martin & Deyrup-Olsen, 1986). In these species, defence adhesives typically gain their mechanical strength from a network of proteins and polysaccharides that stiffen in the presence of substantial metal-binding proteins by forming a network of cross-linked proteins (Braun, Menges, Opoku, & Smith, 2013; Pawlicki et al., 2004; Smith, 2002; Smith, 2006; Werneke, Swann, Farquharson, Hamilton, & Smith, 2007). These products are released from mucus glands in the dorsum as microscopic packets that rupture by ATP or shear stress to form a uniform viscoelastic secretion (Luchtel et al., 1991; Smith, 2010; Werneke et al., 2007). Without sheer, the mucus would not possess adhesive properties and it would just flow off the slug (Deyrup-Olsen et al., 1983).

The ability of mucus packets to rupture in the presence of stress suggests that rubbing of the dorsal surface may trigger the formation of mucus, which would account for the initiation of secretion in *T. graeffei* upon tactile stimulation, as well as the specificity of mucus secretion to the specific dorsal regions disturbed. It is also possible that stimulation of the dorsum leads to contractions within the immediate area, which may be the primary mechanism used for the expression of the mucus to the surface, though this process requires further investigation. Another observation is that the red mucus expressed during periods of stimulation appears to be derived from the red epidermis that skirts the perimeter of the slug’s foot, with cells from this section possibly becoming dislodged into the translucent mucus when the epidermis in this region is damaged. Although this particular mucus didn’t seem to have adhesive properties of its own, further research may determine if it has other important properties such as making the slug unpalatable or reacting with the adhesive secretions to increase its adhesiveness and effectiveness. Nevertheless, this defence mechanism in *T. graeffei* is likely to differ from previously studied species. As such, there is potential for comparative studies to be conducted, since the biochemical makeup and secretory structures vary amongst gastropod species (Foltan, 2004; Smith, 2010).

A strong bioadhesive is a valuable defensive tool for a terrestrial slug that is slow and often chooses to move when conditions are moist. However, this strategy is likely to come at a considerable cost to the slug in terms of future survival, as mucus production requires an investment of water that usually derives from the animal itself. Like many other biological adhesives, the mucus of *T. graeffei* is able to adhere strongly and non-specifically which, along with its ability to maintain or even strengthen its hold under wet conditions, makes it potentially useful in the development of new generations of glues (Callow & Callow, 2006), such as the recent medical adhesives developed based on studies of *A. subfuscus* secretions. Furthermore, investigating the mucus packet systems in this and other species may also help in understanding their ability to quickly harden the mucus secretions, which could provide guidance for future designs in fast-reacting artificial systems.

Adaptations that allow species to defend themselves from predatory attack, particularly those species lacking in speed or camouflage, are evident throughout the animal kingdom. Although not common, the use of adhesives to avoid predation has been found in many types of animals, and typically assumed to make the prey unpalatable or difficult to consume. The ability to use adhesives to immobilize predators has only been examined in the laboratory on amphibians (Arnold, 1982; Evans & Brodie, 1994) and arthropods (Betz & Kölsch, 2004), and observations made on hagfishes which use adhesive mucus to suffocate predators by clogging their gills (Zintzen et al., 2011). However, this appears to not only be the first description of defensive mucus production in this slug species but one of the first natural observations of the use of adhesive defence secretions incapacitating a predator. As such, this finding is important from a natural history perspective, especially since the evolution of defensive mucus secretions in most terrestrial slugs remains unknown. Detailed research on the biochemical structure of such bioadhesives may lead to significant advancements in the development of new materials that are able to exploit their unique mechanical properties.

## Acknowledgements

The authors thank William Legge for providing technical support during fieldwork. This work was conducted under ethics number A-2011-138 approved by the University of Newcastle. All experimental procedures were performed in accordance with the Australian code for the care and use of animals for scientific purposes.

## References

Arnold, S. J. (1982). A Quantitative Approach to Antipredator Performance: Salamander Defense against Snake Attack. Copeia, 1982(2), 247–253. doi:10.2307/1444602

Baer, A., & Mayer, G. (2012). Comparative anatomy of slime glands in Onychophora (velvet worms). Journal of morphology, 273(10), 1079–1088. doi:10.1002/jmor.20044

Barber, J. R., Leavell, B. C., Keener, A. L., Breinholt, J. W., Chadwell, B. A., McClure, C. J., … Kawahara, A. Y. (2015). Moth tails divert bat attack: evolution of acoustic deflection. Proceedings of the National Academy of Sciences, 112(9), 2812–2816. doi:10.1073/pnas.1421926112

Betz, O., & Kölsch, G. (2004). The role of adhesion in prey capture and predator defence in arthropods. Arthropod Structure & Development, 33(1), 3–30. doi:https://doi.org/10.1016/j.asd.2003.10.002

Brau, F., Lanterbecq, D., Zghikh, L.-N., Bels, V., & Damman, P. (2016). Dynamics of prey prehension by chameleons through viscous adhesion. Nature Physics, 12, 931. doi:10.1038/nphys3795

Braun, M., Menges, M., Opoku, F., & Smith, A. M. (2013). The relative contribution of calcium, zinc and oxidation-based cross-links to the stiffness of Arion subfuscus glue. Journal of experimental biology, 216(8), 1475–1483.

Callow, J. A., & Callow, M. E. (2006). The Ulva spore adhesive system. In Biological adhesives (pp. 63–78): Springer.

Cook, A. (1987). Functional aspects of the mucus□producing glands of the Systellommatophoran slug, *Veronicella floridana*. Journal of Zoology, 211(2), 291–305.

Deyrup-Olsen, I., Luchtel, D., & Martin, A. (1983). Components of mucus of terrestrial slugs (Gastropoda). American Journal of Physiology-Regulatory, Integrative and Comparative Physiology, 245(3), R448–R452.

Evans, C. M., & Brodie, E. D. (1994). Adhesive Strength of Amphibian Skin Secretions. Journal of Herpetology, 28(4), 499–502. doi:10.2307/1564965

Flammang, P., Demeuldre, M., Hennebert, E., & Santos, R. (2016). Adhesive secretions in echinoderms: a review. In A. M. Smith (Ed.), Biological Adhesives (2nd Edition ed., pp. 193–222). Switzerland: Springer.

Foltan, P. (2004). Influence of slug defence mechanisms on the prey preferences of the carabid predator Pterostichus melanarius (Coleoptera: Carabidae). European Journal of Entomology, 101(3), 359–364.

Graham, L. D., Glattauer, V., Li, D., Tyler, M. J., & Ramshaw, J. A. (2013). The adhesive skin exudate of Notaden bennetti frogs (Anura: *Limnodynastidae*) has similarities to the prey capture glue of Euperipatoides sp. velvet worms (Onychophora: *Peripatopsidae*). Comparative Biochemistry and Physiology Part B: Biochemistry and Molecular Biology, 165(4), 250–259.

Landauer, M. R., & Chapnick, S. D. (1981). Responses of Terrestrial Slugs to Secretions of Stressed Conspecifics. Psychological Reports, 49(2), 617–618. doi:10.2466/pr0.1981.49.2.617

Li, J., Celiz, A. D., Yang, J., Yang, Q., Wamala, I., Whyte, W., … Mooney, D. J. (2017). Tough adhesives for diverse wet surfaces. Science, 357(6349), 378–381. doi:10.1126/science.aah6362

Luchtel, D., Deyrup-Olsen, I., & Martin, A. (1991). Ultrastructure and lysis of mucin-containing granules in epidermal secretions of the terrestrial slug Ariolimax columbianus (Mollusca: Gastropoda: Pulmonata). Cell and tissue research, 266(2), 375–383.

Mair, J., & Port, G. R. (2002). The influence of mucus production by the slug, Deroceras reticulatum, on predation by *Pterostichus madidus* and *Nebria brevicollis* (Coleoptera: Carabidae). Biocontrol Science and Technology, 12(3), 325–335.

Martin, A. W., & Deyrup-Olsen, I. (1986). Function of the epithelial channel cells of the body wall of a terrestrial slug, *Ariolimax columbianus*. Journal of experimental biology, 121(1), 301–314.

Pawlicki, J., Pease, L., Pierce, C., Startz, T., Zhang, Y., & Smith, A. (2004). The effect of molluscan glue proteins on gel mechanics. Journal of experimental biology, 207(7), 1127–1135.

Rice, S. H. (1985). An anti-predator chemical defense of the marine pulmonate gastropod *Trimusculus reticulatus* (Sowerby). Journal of experimental marine biology and ecology, 93(1-2), 83–89.

Rollo, C. D., & Wellington, W. G. (1979). Intra-and inter-specific agonistic behavior among terrestrial slugs (Pulmonata: Stylommatophora). Canadian Journal of Zoology, 57(4), 846– 855.

Schubert, M., Munday, P. L., Caley, M. J., Jones, G. P., & Llewellyn, L. E. (2003). The toxicity of skin secretions from coral-dwelling gobies and their potential role as a predator deterrent. Environmental Biology of Fishes, 67(4), 359–367.

Shephard, K. L. (1994). Functions for fish mucus. Reviews in fish biology and fisheries, 4(4), 401–429.

Smith, A. M. (2002). The structure and function of adhesive gels from invertebrates. Integrative and Comparative Biology, 42(6), 1164–1171.

Smith, A. M. (2006). The biochemistry and mechanics of gastropod adhesive gels. In S. A. M. & C. J. A. (Eds.), Biological Adhesives (pp. 167–182). Berlin, Germany: Springer.

Smith, A. M. (2010). Gastropod Secretory Glands and Adhesive Gels. In J. von Byern & I. Grunwald (Eds.), Biological Adhesive Systems: From Nature to Technical and Medical Application (pp. 41–51). Vienna: Springer Vienna.

Smith, A. M., & Callow, J. A. (2006). Biological adhesives. Berlin, Germany: Springer.

South, A. (2012). Terrestrial slugs: biology, ecology and control. London: Springer Science & Business Media.

Triebskorn, R., & Ebert, D. (1989). The importance of mucus production in slugs’ reaction to molluscicides and the impact of molluscicides on the mucus producing system. University of Tübingen, Heidelberg, Germany.

Verdugo, P. (1991). Mucin Exocytosis1-3. The American Review of Respiratory Disease, 144, S33– S37.

von Byern, J., Müller, C., Voigtländer, K., Dorrer, V., Marchetti-Deschmann, M., Flammang, P., & Mayer, G. (2017). Examples of Bioadhesives for Defence and Predation. In S. N. Gorb & E. V. Gorb (Eds.), Functional Surfaces in Biology III: Diversity of the Physical Phenomena (pp. 141–191). Cham: Springer International Publishing.

Werneke, S., Swann, C., Farquharson, L., Hamilton, K., & Smith, A. (2007). The role of metals in molluscan adhesive gels. Journal of experimental biology, 210(12), 2137–2145. doi:10.1242/jeb.006098

Wilks, A. M., Rabice, S. R., Garbacz, H. S., Harro, C. C., & Smith, A. M. (2015). Double network gels and the toughness of terrestrial slug glue. Journal of experimental biology, jeb. 128991.

Zintzen, V., Roberts, C. D., Anderson, M. J., Stewart, A. L., Struthers, C. D., & Harvey, E. S. (2011). Hagfish predatory behaviour and slime defence mechanism. Scientific Reports, 1, 131. doi:10.1038/srep00131

